# Dynamic Sex Chromosome Expression in Drosophila Male Germ Cells

**DOI:** 10.1101/2020.03.23.000356

**Authors:** Sharvani Mahadevaraju, Justin M. Fear, Miriam Akeju, Brian J. Galletta, Mara MLS. Pinheiro, Camila C. Avelino, Diogo C. Cabral-de-Mello, Katie Conlon, Stafania Dell’Orso, Zelalem Demere, Kush Mansuria, Carolina A. Mendonça, Octavio M. Palacios-Gimenez, Eli Ross, Max Savery, Kevin Yu, Harold E. Smith, Vittorio Sartorelli, Nasser M. Rusan, Maria D. Vibranovski, Erika Matunis, Brian Oliver

**Affiliations:** Laboratory of Cellular and Developmental Biology, National Institute of Diabetes and Kidney and Digestive Diseases, National Institutes of Health, Bethesda, MD 20892, USA; Department of Cell Biology, Johns Hopkins University School of Medicine, 725 N. Wolfe Street, Baltimore, MD 21205, USA; Cell Biology and Physiology Center, National Heart, Lung and Blood Institute, National Institutes of Health, Bethesda, MD 20892, USA; Department of Genetics and Evolutionary Biology, Institute of Biosciences, University of São Paulo, SP 05508-090, São Paulo, BRA; Instituto de Biociências/IB, Departamento de Biologia Geral e Aplicada, UNESP— Universidade Estadual Paulista, Rio Claro, São Paulo 13506-900, BRA; Laboratory of Muscle Stem Cells and Gene Regulation, National Institute of Arthritis and Musculoskeletal and Skin Diseases, National Institutes of Health, Bethesda, MD 20892, USA; Department of Evolutionary Biology, Evolutionary Biology Centre, Uppsala University, 75236, Uppsala, SE; Department of Organismal Biology, Systematic Biology, Evolutionary Biology Centre, Uppsala University, 75236, Uppsala, SE; Genomics Core, National Institute of Diabetes and Kidney and Digestive Diseases, National Institutes of Health, Bethesda, MD 20892, USA

## Abstract

Sex chromosome gene content and expression is unusual. In many organisms the X and Y chromosomes are inactivated in spermatocytes, possibly as a defense mechanism against insertions into unpaired chromatin. In addition to current sex chromosomes, Drosophila has a small gene-poor X-chromosome relic (4^th^) that re-acquired autosomal status. Using single cell RNA-Seq, we demonstrate that the single X and pair of 4^th^ chromosomes are specifically inactivated in primary spermatocytes. In contrast, genes on the single Y chromosome become maximally active in primary spermatocytes. Reduced X steady-state transcript levels are due to failed activation of RNA-Polymerase-II by phosphorylation of Serine 2 and 5.

**One Sentence Summary:** Sex chromosome expression during spermatogenesis at the single cell level

## Main Text

Organisms as diverse as flies and humans have XY males and XX females. In Drosophila, the large gene-poor Y chromosome, inherited solely from males, is heterochromatin-rich and carries a few genes essential for fertility (*1, 2*). The X chromosome is present as a single copy in males. The 4^th^ chromosome is an ancestral X chromosome (*3*) and shares some aspects of sex chromosome structure, such as being heterochromatin-rich (*4*). Given the vast difference between sex chromosomes and their unique modes of inheritance, the unusual gene content and expression of sex chromosomes is widely studied (*5*). In many organisms, including mammals (*6*), *C. elegans (7)*, and Drosophila (*8*), X expression is reduced in testis. Non-mutually exclusive reasons for this reduction include: inactivation of unpaired chromosomes (*9, 10*), absence of germline X chromosome dosage compensation (*11*), and evolutionary re-localization of genes required in males off the X chromosome (*8, 12, 13*).

Meiotic X-chromosome inactivation is well described in mammals, where premature transcriptional inactivation of the X and Y chromosome during mid-spermatogenesis is essential for fertility (*14*). The X and Y preciously condense into a heterochromatic “XY body” within the pachytene spermatocyte nucleus. Transcriptional inactivation is thought to protect these largely dissimilar chromosomes by defending un-synapsed regions of the genome from stealthy invasive transposons that cannot be detected as foreign in the absence of a homolog (*15*). Inactivation may also protect against unwanted recombination between the X and Y, or to repair damage created by lack of recombination (*16*). Inactivation might also mark paternal X chromosomes for later imprinting for inactivation (*17, 18*). Some of these models are unlikely in the case of Drosophila males, which, for example, completely lack meiotic recombination (*19*). Exploiting differences, and commonalities, between Drosophila and other models can illuminate causes of unusual sex chromosome expression patterns.

In many organisms, the potentially lethal imbalance of X-linked gene transcripts from the single X relative to the paired autosomes must be alleviated in the soma to ensure viability (*20*). In Drosophila somatic cells, this X chromosome dosage compensation is also essential and involves large-scale transcriptional up-regulation of the single X chromosome (*21*). It is unclear if dosage compensation exists in Drosophila germ cells, which do not require the core dosage compensation machinery (*22, 23*) suggesting that either dosage compensation does not occur, or if it does, that it relies on a novel mechanism. Analysis of transcripts expressed in portions of hand-dissected testes (*24, 25*) or in mutants that block spermatogenesis in mitotic stages (*13, 20, 26*) suggest that Drosophila testis enriched in mitotic spermatocytes show at least partial X chromosome dosage compensation. Portions of adult testes enriched in meiotic spermatocytes (*24*) show greatly reduced X chromosome expression and autosomal genes inserted on the X show dramatically reduced expression in spermatocytes, far exceeding simple failure of dosage compensation (*22, 23, 27*), consistent with X inactivation. Given the potentially dynamic expression of the X chromosome by stage and cell type, cellular resolution of sex chromosome expression during spermatogenesis is needed. We did this by single cell RNA sequencing (scRNA-Seq) (*28*).

Drosophila third instar larval (L3) testis contain abundant germ cells, including the critical transition from mitotic spermatogonia to meiotic primary spermatocytes (*29, 30*). Prior to using L3 testes for single cell analysis, we asked whether the patterns of chromosome-specific gene expression known to occur in adult gonads (*8*) are also present at this earlier stage of gonadogenesis. We performed RNA-Seq on eight biological replicates each of cleaned 3^rd^ instar larval testes and ovaries (**Fig. S1)** to compare with recent adult profiles from our laboratory (*31*). The overall levels of X-linked gene expression from the single X in testis was reduced in both adult and larva (**Fig. 1A, B**). Interestingly, the two 4^th^ chromosomes, derived from an ancient X chromosome that re-acquired autosome status (*3*), also showed a dramatic decrease in overall expression in testis relative to ovary. Expression of the male-specific Y chromosome was highly testis-biased. Like adult testis, larval testis exhibit unusual sex chromosome expression patterns.

**Figure 1.**
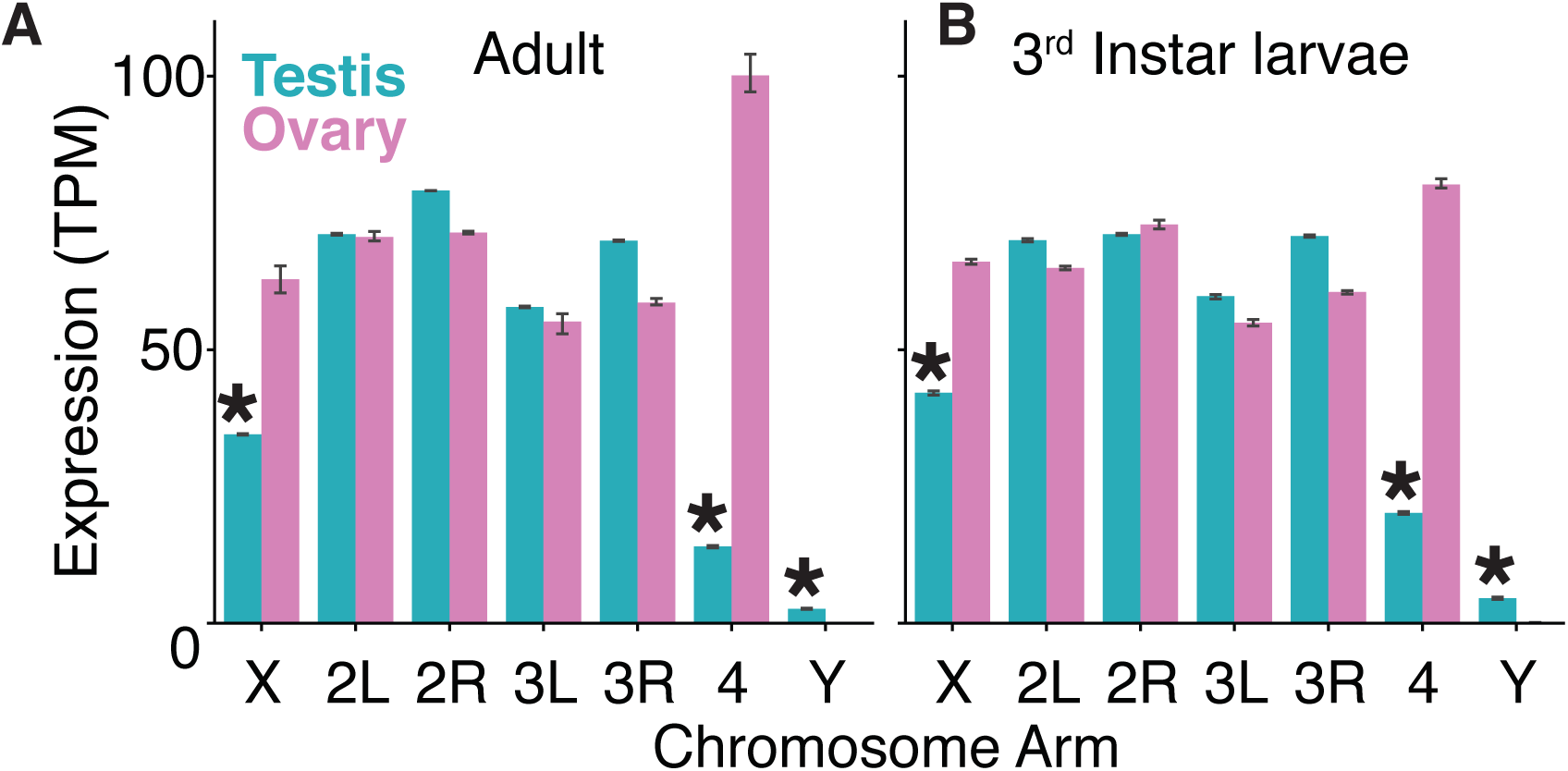
Whole-tissue RNA-Seq of adult and larval gonads. Comparison of gene expression by chromosome arms using the average number of RNA-Seq reads, normalized for gene length and library size (Transcripts Per Million reads = TPM) mapping to each chromosome arm from testis (blue) and ovary (pink) libraries for (**A**) adults and (**B**) 3^rd^ instar larvae. Bars are 95% confidence intervals of the mean as determined by bootstrapping. Comparison of testis versus ovary P ≤ 0.01, one-sided Welch’s t-test (*).

Determining which cells of the testis contribute to unusual patterns of sex chromosome expression is challenging because of the cellular diversity and dynamic nature of gene expression in the gonad. Drosophila spermatogenesis (**Fig. 2A**) begins at the apical end of the testis (*32*) where germline stem cells divide asymmetrically, generating mitotically active differentiating daughters called gonialblasts that produce clusters of interconnected spermatogonia (G). After their fourth mitotic division, spermatogonia quickly enter pre-meiotic S-phase, becoming early primary spermatocytes (E1°) that are initially morphologically indistinguishable from spermatogonia (*33*). As early primary spermatocytes traverse an extended G2 phase, they undergo a burst of transcriptional activity that is accompanied by a 25-fold increase in volume (*30*), becoming large middle primary spermatocytes (M1°) and then slightly smaller late primary spermatocytes (L1°). Fully mature late primary spermatocytes progress through two rapid meiotic divisions, becoming spermatids. Pairs of quiescent somatic cyst cells (C1-C4) envelope each gonialblast and descendants, differentiating alongside the germ cells they support. Cyst cells are a source of intercellular signals and act as a permeability barrier (*34, 35*). The entire L3 testis is encapsulated by an epithelial monolayer of pigment cell precursors (P), which become adult pigment cells and are required for joining the gonad with the reproductive tract (*36*). The basal end of the L3 testis contains terminal epithelial precursor cells (T) which will ultimately regulate the final steps of spermiogenesis (*37*).

**Figure 2.**
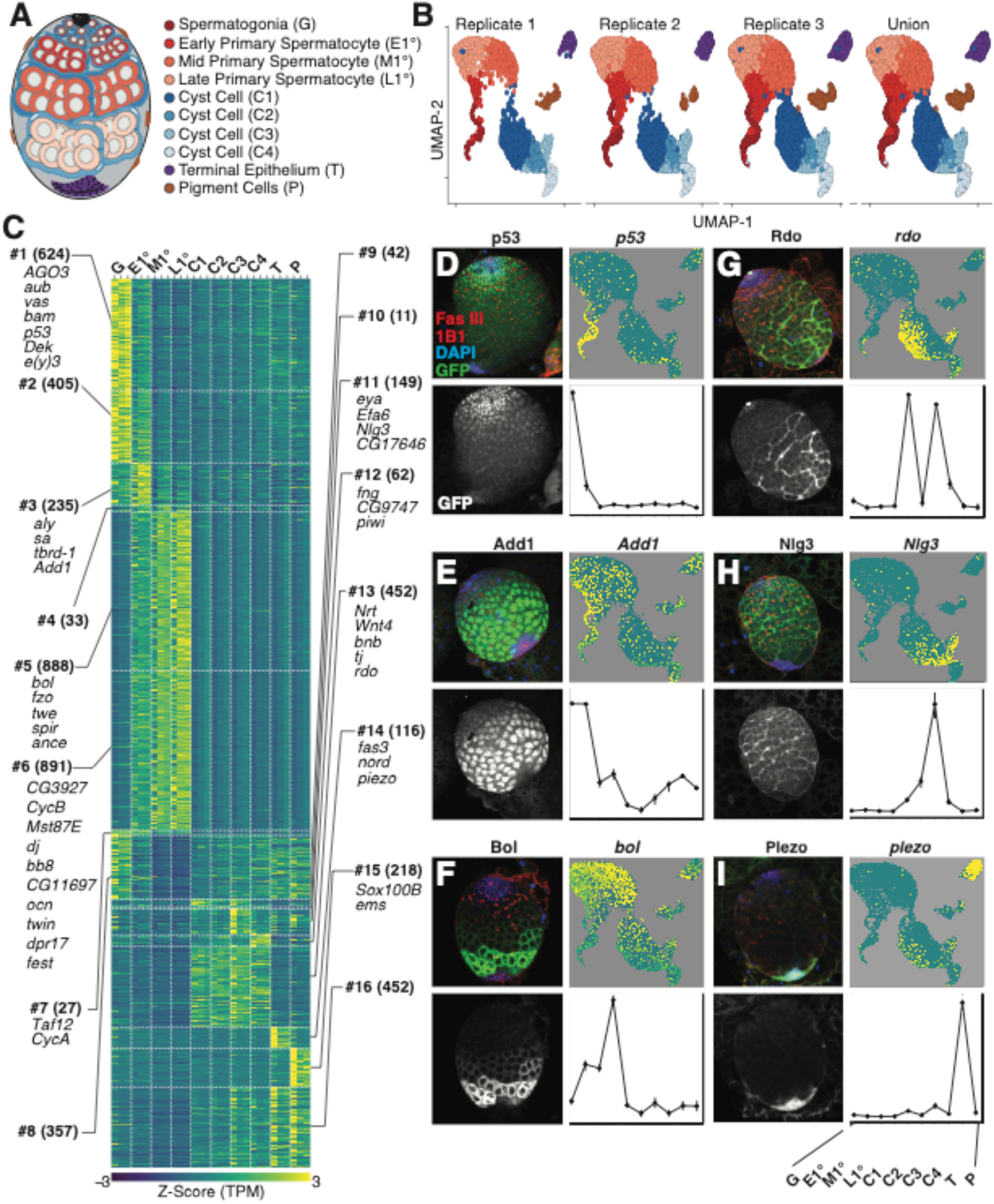
Identification and annotation of larval testis cell types from scRNA-Seq. (**A**) Illustration of the 3^rd^ instar larval testis. Abbreviations and color code for each cell type are shown. (**B)** Uniform Manifold Approximation and Projection (UMAP) for each scRNA-Seq replicate and the union of all three data sets. Each cell is projected onto the two-dimensional UMAP space and clustered using k-nearest neighbors. Cell-types are colored to correspond to the testis model. **(C)** Patterns of differential gene expression among putative cell types. Gene expression (Z-score; low: blue, high: yellow) for each differentially expressed gene (rows) for all cell-type-by-replicate (columns). Genes are ordered and grouped (horizontal dashed lines) manually into 16 classes (numbered label) based on differential gene expression. Each class is annotated with the number of genes (in parentheses) and the key genes used to identify the cell types (**Table S2**). (**D-I**) Immunofluorescence images showing protein expression of representative genes from the protein trap reporters for cell type annotation. (**D**) *p53*, **(E)** *Add1* (**F**) *bol* (**G**) *rdo* (**H**) *Nlg3*, and (**I**) *Piezo*. Each panel consists of color (Fas III, 1B1: red, DAPI: blue, and GFP: green) and greyscale (GFP) images of protein trap expression, scRNA-Seq gene expression projected onto the UMAP (Z-score; low: blue, high: yellow), and normalized gene expression values plotted by cell type.

To capture the transcriptional profiles of the cell types in the Drosophila L3 testis, we dissected staged male third instar larvae, enzymatically removing the associated fat body before dissociating 20-40 testes to a single cell suspension for scRNA-Seq (**Materials and Methods, Fig. S1A**-**D**). We piloted dissociation conditions using fly strains expressing GFP in subsets of cells (**Fig. S1G-H**) coupled with dye exclusion (**Fig. S1I**) (70-75% viability, n =1,000/replicate), ensuring consistent retrieval of single cell suspensions of viable germline and somatic cells. We then performed scRNA-Seq on wild-type L3 testes. We identified 18,965 single cells across three biological replicates (Spearman ρ ≥ 0.93, P < 0.001; **Table S1**) based on the intersection of calls from *cell ranger count (38)* and *DropletUtils emptyDrops (39)* (**Table S2**). Potential cell doublets were detected using *scrublet* (*40*) and removed. We observed a significant match of the sum of scRNA-Seq cell-based reads with replicated RNA-Seq from whole L3 testes (Spearman ϱ ≥ 0.77, P < 0.001; **Table S1**), indicating that major cell types are well represented in our scRNA-Seq dataset. Based on preliminary cluster analysis, we set the perplexity threshold in *Seurat* to 0.3 using the 2,000 most variably expressed genes. This yielded ten clusters, each potentially representing a distinct cell type or state. We obtained similar cluster profiles from individual scRNA-Seq replicates as shown by Uniform Manifold Approximation and Projection (UMAP), showing concordance across the three biological replicates (**Fig. 2B**). Count tables showing all data (**Table S2**) and UMAP projections for each gene are available (**Fig. S2**).

We identified genes with enriched expression in cell types by comparing the expression profiles of cells in each of the ten groups with that of all remaining cells in an iterative process for each cell cluster (Wilcoxon rank-sum test, FDR corrected P ≤ 0.01; **Fig. 2C, Table S2**). To determine what the cell types were, we projected genes with well-known expression patterns, carefully re-curated from published images, onto the cell level expression profiles. We also used transgenic protein-trap reporter genes (*41*) to identify expression patterns for genes with enriched expression in each cell type (**Fig. 2D-I, Fig. S3, Table S3)**. This allowed us to annotate all ten clusters. 91% of 79 genes from the literature or reporter expression patterns overlapped cell types identified by scRNA-Seq. 46% were expressed in the same or developmentally preceding cell types, and 45% were expressed in the same cell lineage in both the scRNA-Seq clusters and imaged testis. The 7% of genes showing discordant assay-dependent expression patterns were protein-trap reporters showing either weak or widespread expression, both of which hinder assignment to specific cell types (**Fig. S3, Table S3**). These data indicate that we have identified many of the major cell types in the L3 testis. We did not identify new cell types and failed to unambiguously identify cells in the stem cell niche. A few examples of the vast number of genes with interesting expression profiles follow.

Spermatogonia (G) were characterized by the biased expression of 624 genes (**Fig. 2C #1)** including: known early germ cell markers *argonaute 3* (*AGO3*), *aubergine* (*aub*), and *vasa* (*vas*) (*42*), the early differentiation signal *bag-of-marbles* (*bam*) (*33*), the mitotic cell marker *p53 (43)*(**Fig. 2D**), the chromatin proteins *Dek* (*44*) and *enhancer of yellow 3* (*e(y)3* (*45*) (**Fig. 2C #1)**. While expression of these spermatogonial genes also extended to spermatocytes, we could readily distinguish spermatogonia from spermatocytes by their lack of spermatocyte-specific gene expression. The earliest primary spermatocytes (E1°) were characterized by high expression of 235 genes including three markers of the transition from spermatogonia to spermatocyte: *always early* (*aly*), *spermatocyte arrest (sa)* (*46*), and *testis-specifically expressed bromodomain containing protein-1* (*tbrd-1*) (*47*) (**Fig. 2C #3**). Sa is a testis-specific TATA-Binding Protein Associated Factor (tTAF), Aly serves as a testis-specific meiotic arrest complex (tMAC) member (*46*), and tbrd-1 is associated with both transcriptional complexes (*48*) (**Fig. 2E**). In addition, we show that *ADD domain-containing protein 1* (*Add1)*, which encodes a heterochromatin associated protein that interacts with Heterochromatin Protein 1 (HP1) to maintain heterochromatin (*49, 50*), has enriched expression in primary spermatocytes. The tTAFS and tMACs initiate expression of a large battery of genes, many expressed exclusively in primary spermatocytes (*51*). Middle and late primary spermatocytes (M1^0^ and L1°) are characterized by the increased expression of two large groups of 1,779 genes that are targets of tTAFs and tMACs (**Fig. 2C #5, #6)**, including *don juan* (*dj*) (*52*). Interestingly, *ocnus* (*ocn)* transgenes show very poor expression when inserted on the X chromosome versus when inserted on autosomes suggesting that *ocn* reports X inactivation in Drosophila (*27*). As *ocn* is expressed in primary spermatocytes (**Fig. 2C #6)**, this suggests that inactivation occurs in primary spermatocytes. As spermatocytes progress through differentiation, we observed a concomitant increase in the expression of the meiotic cell cycle regulator *twine* (*twe*) and the translational regulator of Twe encoded by *boule* (*bol)* (*53, 54*) by both scRNA-Seq and protein-trap analysis (**Fig. 2F)**. These data defined four distinct stages in male germ cell development in larval testis.

The somatic cell types provide an important baseline for analysis of germline expression and contribute to further understanding of these cell types in the testis (**Fig. 2C**). For example, *traffic jam* (*tj*) and *eyes absent (eya)* are expressed in early and late cyst cells respectively in both adult and L3 testis (*55, 56*), and are most highly expressed in distinct cyst cell clusters in this study. There is also a great deal of new biology in the cell expression profiles. We discovered that the transmembrane protein encoded by *defective proboscis extension response 17* (*dpr17*), a neuronal surface label, is enriched in spermatocytes. We also discovered that the genes encoding the visual system Reduced ocelli (Rdo) protein (*57*) (**Fig. 2G**), the exchange factor for the GTPase Arf-6 (Efa6) (*58*), and the synaptic adhesion molecule Neuroligin 3 (Nlg3) (*59*) (**Fig. 2H**), show dynamic expression patterns in the cyst cells. The terminal epithelium is poorly studied and we report multiple new RNA and protein-trap markers for this cell type, including the human neuron derived neurotrophic factor (Nord) homolog, as well as the mechanosensory ion channel subunit Piezo *(60)* (**Fig. 2I**), which acts as a stretch sensor and could help regulate transit of sperm from the testis into the seminal vesicle. This data should be an outstanding resource for those studying testis development and physiology.

We looked at the dynamics of sex chromosome gene expression in germ cells in addition to all the other cell types from the single cell dataset. Since genes with high expression in the testis are not uniformly distributed in the genome (*8, 13*), to avoid the confusion that the reduced expression from the X could be because of reduced density of testis-biased genes we assayed different gene sets. We analyzed “all expressed genes”, a set of 14,347 genes expressed in at least one cell type and “widely expressed genes”, a set of 589 genes expressed in ≥ 33% of all cells in the single cell data. The “Housekeeping genes”, a name given to a set of genes based on expression in a wide range of Drosophila tissue (tau and TSPS) (*61, 62*) is inappropriate for our analysis as it showed poor expression in germ cells.

Expression of the single X chromosome relative to the major autosomes (chromosomes 2 and 3, each present in two copies) is not significantly different in spermatogonia or any of the somatic cell types using either all expressed genes or widely expressed genes (**Fig 3A, B**). The somatic cell expression pattern is consistent with the known canonical X chromosome dosage compensation mechanism in somatic cells (*21*). Our data provides new evidence for non-canonical dosage compensation of the X chromosome in spermatogonia. Interestingly, at least some genes on mammalian X chromosomes are also overexpressed prior to inactivation (*63*), suggesting that this dynamic transition in X chromosome expression in the male germline is conserved. There was a significant and progressive decrease (P ≤ 0.001) in steady-state expression of the X chromosome in early, middle and late primary spermatocytes (E1°, M1°, and L1°). This decrease in X expression could be due to either a loss of dosage compensation in germ cells as they mature into primary spermatocytes, or to the gain of meiotic X-chromosome inactivation, as seen in mammalian primary spermatocytes. Expression of 4^th^ chromosome genes paralleled what was seen for the X. There was a significant and progressive decrease (P < 0.001) in steady-state expression levels in M1° and L1° (**Fig. 3C, D)** compared to expression in spermatogonia. Because all autosomes, including the 4^th^, are present in two copies in both males and females, this decreased expression of 4^th^ chromosome genes cannot be due to loss of dosage compensation. Instead, a gain of inactivation is the simplest explanation. Finally, the entire pattern of Y chromosome expression was inverted relative to that seen from the X and 4^th^ chromosomes (**Fig. 3E**). We observed poor expression of the Y in somatic cells, and increased expression in E1°, M1°, and L1° primary spermatocytes. This is likely to occur from expression of a few highly transcriptionally active Y-linked genes originally identified by the cytologically visible Y-chromosome loops present at these stages (*64, 65*). The decrease in sex chromosome expression in M1° and L1° did not reflect an overall decrease in total gene expression compared to somatic lineages (**Fig. 3F**). Since the single X and the two 4^th^ chromosomes showed reduced expression, while the single Y showed ongoing expression, these data demonstrate that there is no simple rule for sex chromosome gene expression in Drosophila primary spermatocytes.

**Figure 3.**
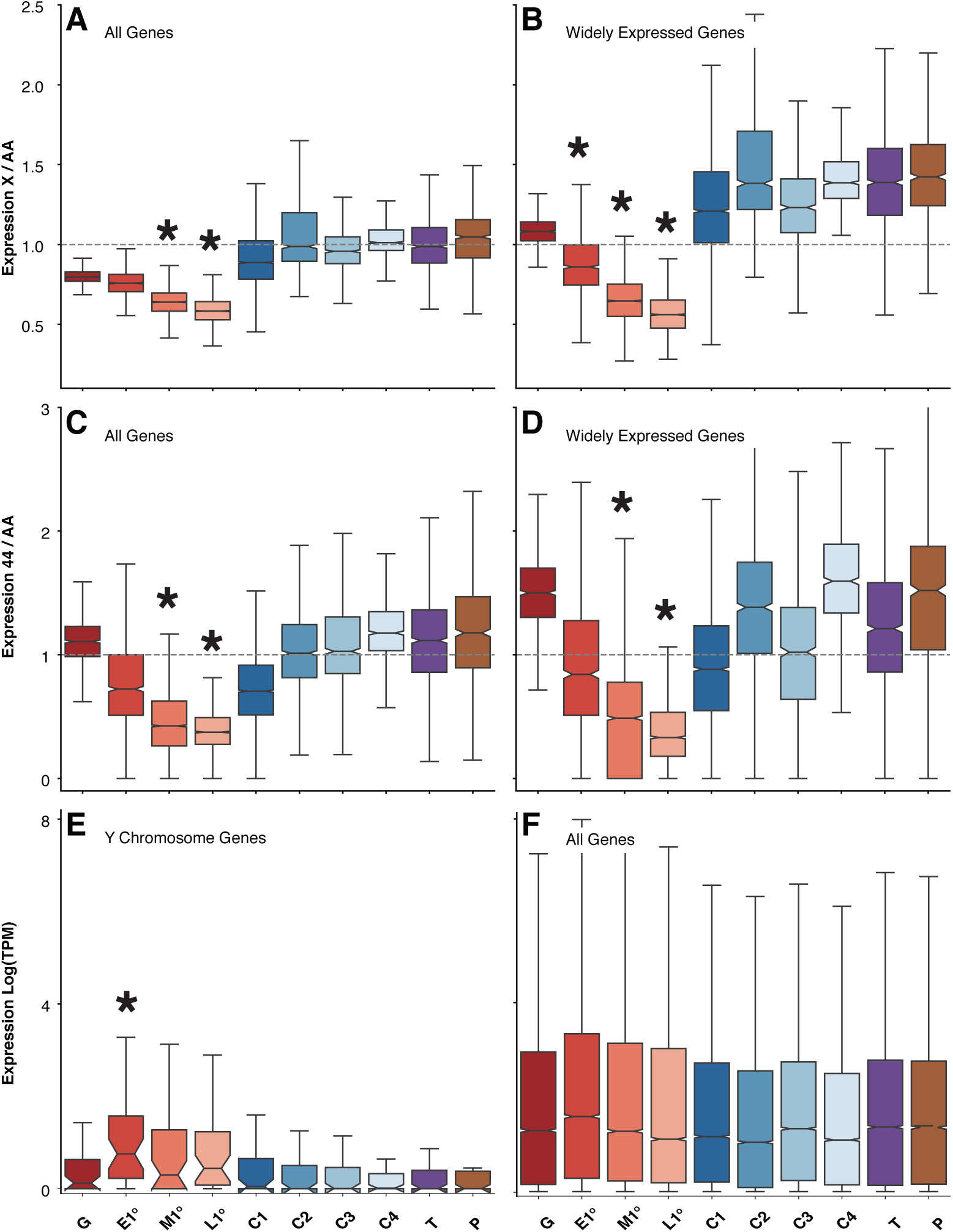
X-, 4^th^-, and Y-linked gene expression during spermatogenesis. Cell type-specific gene expression of X-linked genes (**A-B**) and 4^th^-linked genes (**C-D**) relative to the major 2^nd^ and 3^rd^ chromosomes. Y-linked gene expression in normalized read counts ln(TPM + 1) (**E**), and genome-wide expression in normalized read counts ln(TPM + 1) (**F**). Chromosome-specific gene expression is relative to all expressed genes in the data set (**A, C**, and **G**), or a subset of genes widely expressed across cell types (**B** and **D**). X-linked (**A-B**) and 4^th^-linked (**C-D**) gene expression are scaled by major autosomal arm gene expression. Chromosomal distributions in germ cells, significantly different from somatic cell types, are indicated with * (**A-D**: *P* ≤ 0.01 using permutation test, **E:** *q*-value ≤ 0.01 using Chi-Square test). (**A-F**) Boxplots (box = interquartile range (IQR), notch = 95% confidence interval of median, whiskers = ±1.5xIQR).

Given that the steady state level of X expression falls in maturing primary spermatocytes, and since poorly expressed regions of the genome are generally heterochromatic, we sought an association of expression with chromosome state in primary spermatocytes. To locate chromosomes, we used satellite and oligopaint probes that were validated on somatic metaphase chromosomes (**Fig. S4**). The major autosomal bivalents and the X chromosome reside in three distinct chromosome territories that abut the nuclear envelope of primary spermatocytes (*66*). The nuclear interior, less densely stained for DAPI, is occupied by the transcriptionally active Y chromosome (*64, 65, 67*) (**Fig. 4A**). *In situ* hybridization reveals X chromatin heterochromatic satellite sequences near the prominent spermatocyte nucleolus, where Ribosomal DNA repeats are located, and ribosome biogenesis occurs (**Fig. 4B**) (*68*). The only homology between the X and Y chromosome is the rDNA clusters. Interestingly, only the Y-linked rDNAs are expressed; the X-linked cluster is inactive (*69*), which is consistent with the pattern we observed chromosome-wide. The probes recognizing satellite sequences of the Y chromosome showed a patchy distribution in the region separating the three main chromosome territories (**Fig. 4B**) (*64*). The 4^th^ is often, but not always, near the nucleolus (*67*). We observed this as well (**Fig. 4C, Fig. S5**).

**Figure 4.**
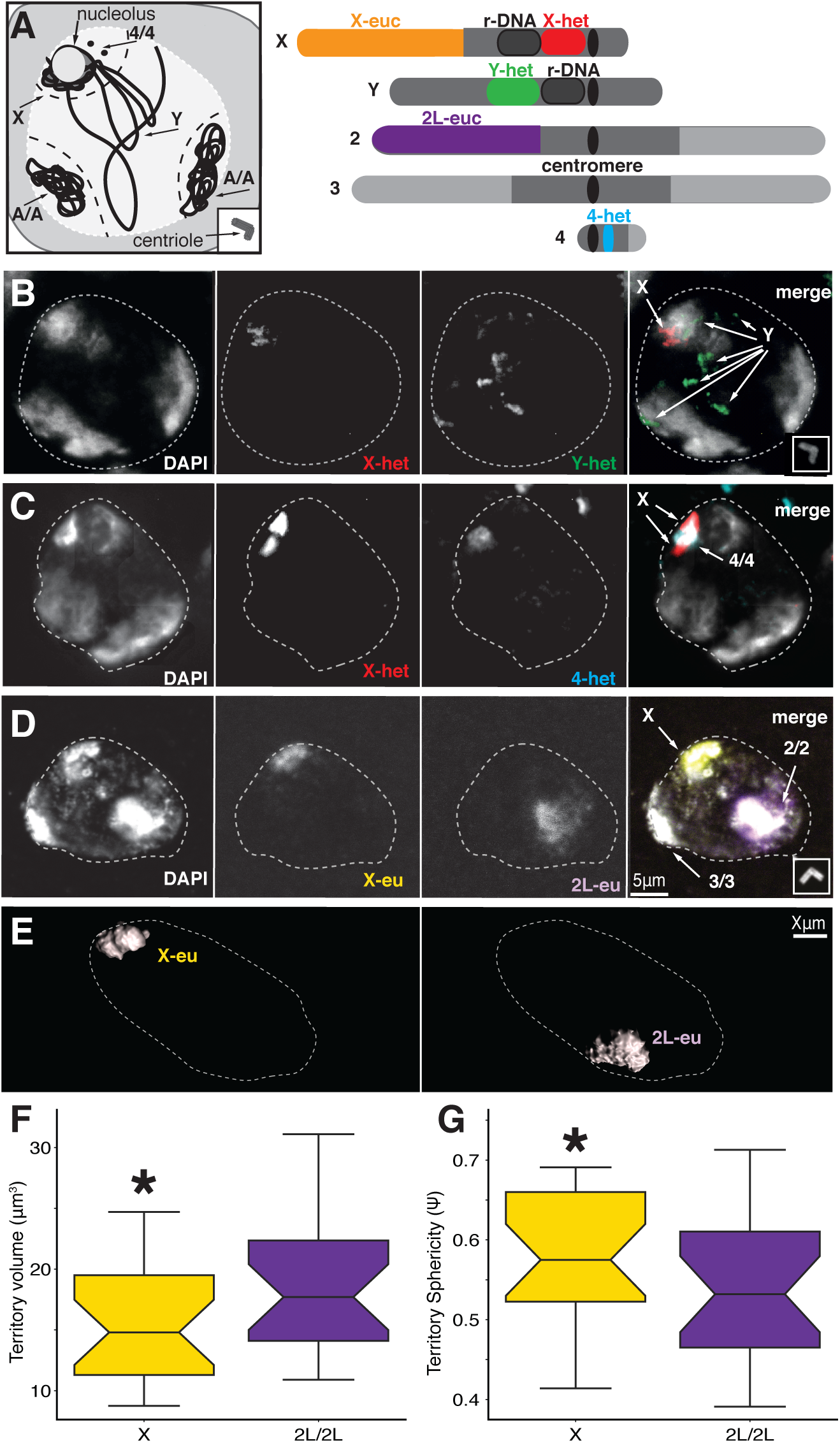
Chromosomal territories in late primary spermatocyte nuclei. (**A**) Cartoon of spermatocyte nuclear structure showing cytoplasm (grey) and nucleus (light grey) with the three chromosome territories indicated (dashed lines) along with locations for the single X and Y, the major autosomes (A/A) and the dot 4th chromosome (4/4) (arrows). The nucleolus (arrow) associates with the X territory. Centriole replication (inset) marks late G2 stages. (**B**) Cartoon of spermatocyte chromosomes. Centromeres (black) heterochromatin (grey), euchromatin (light grey), and rDNA (dark grey) with approximate locations of probes used (color); 1.688 satellite (red), AATAC satellite (green), AATAT satellite (blue), X-euchromatin oligopaints (yellow) and 2L-euchromatin oligopaints (purple). Size not to scale. (**C-E**) channel separated and merged Images of hybridized spermatocyte nuclei. Color coded as in (**B**) with DAPI (grey) and centrioles (insets) stained with anti-Asterless. Primary spermatocyte nucleus is outlined (white-dashed line) based on DAPI and pilot work using anti-laminB. Deduced chromosome identities are shown (arrows). **(F**) Masked rendering of the X and 2L locations and volumes in co-stained nuclei. (**G-H**) boxplots (box = interquartile range (IQR), notch = 95% confidence interval of median, whiskers = ±1.5xIQR) showing (**G**) volume and (**H**) sphericity of the X (yellow) versus 2L (purple). (N = 23). P-value ≤ 0.01, Welch’s t-test (*).

We were most interested in chromosome structure in gene rich euchromatic regions. Hence, we used oligopainting to examine the euchromatic portions of the X chromosome for evidence of compaction that might accompany inactivation. Oligopaint probes show that inactive X chromosomes have increased sphericity and greater compaction than active X chromosomes in mammalian cells (*67, 70*). We probed similar sized euchromatic regions of the X chromosome (22.3 Mb) and the left arm of the 2nd chromosome (2L, 22.7 Mb) with oligopaints (**Fig. 4D**). We converted raw *in situ* data in *Imaris* to create masks (n=23) of pixel intensities (**Movie1**) and obtained volumetric measurements of these territories (**Fig. 4E**). We did find a significant difference between X chromosome and 2L volume when we did not correct for copy number (**Fig 4F**), and the single X (14.8 µm) took a larger volume than half of the two 2L arms (17.7 µm), which is inconsistent with compaction of the X (we do not know if the relationship between volume and copy number is linear). In contrast, the X chromosome was more spherical than 2L (**Fig 4G**), which is consistent with the hypothesis that compaction regulates X expression in primary spermatocytes.

We more directly addressed transcriptional status of chromosome territories in spermatocytes by determining the phosphorylation status of the regulatory C-Terminal Domain (CTD) of RNA Polymerase II (Pol-II). The transcription cycle begins when Pol-II binds the promoter. Serine 5 phosphorylation (pSer5-CTD) causes RNA Pol-II to initiate transcription, and subsequent Serine 2 phosphorylation (pSer2-CTD) induces transcriptional elongation (**Fig. 5A**) (*71*). In the whole testis, we observed pSer2-CTD in spermatogonia and in the two autosomal territories in spermatocytes (**Fig. 5B**). In spermatocyte, we observed clear pSer2-CTD in two autosomal territories, but the X territory (identified by nucleolar proximity) was poorly stained as previously reported (*72*) and pSer5-CTD localization was similar to pSer2-CTD pattern (**Fig. 5C-D)**. We also observed localization of total Pol-II, but very little pSer2-CTD on the X territory when we co-stained with X euchromatic oligopaints (**Fig. S6**), consistent with the idea that reduced steady-state levels of X chromosome transcripts may be due to decreased active transcription. To obtain more quantitative information on transcriptional status of the X chromosome, we determined the ratio of pSer2-CTD and pSer5-CTD to total Pol-II within a given territory compared to the autosomes in late primary spermatocytes (**Fig. 5C-D)**. We found a >2-fold decrease in pSer2-CTD and in pSer5-CTD on the X relative to the autosomes in late primary spermatocytes (**Fig. 5E-F**). These data indicate that the decline in steady-state X chromosome transcripts seen by scRNA-Seq is due to a block in the transcriptional cycle regulated by CTD tail phosphorylation.

**Figure 5.**
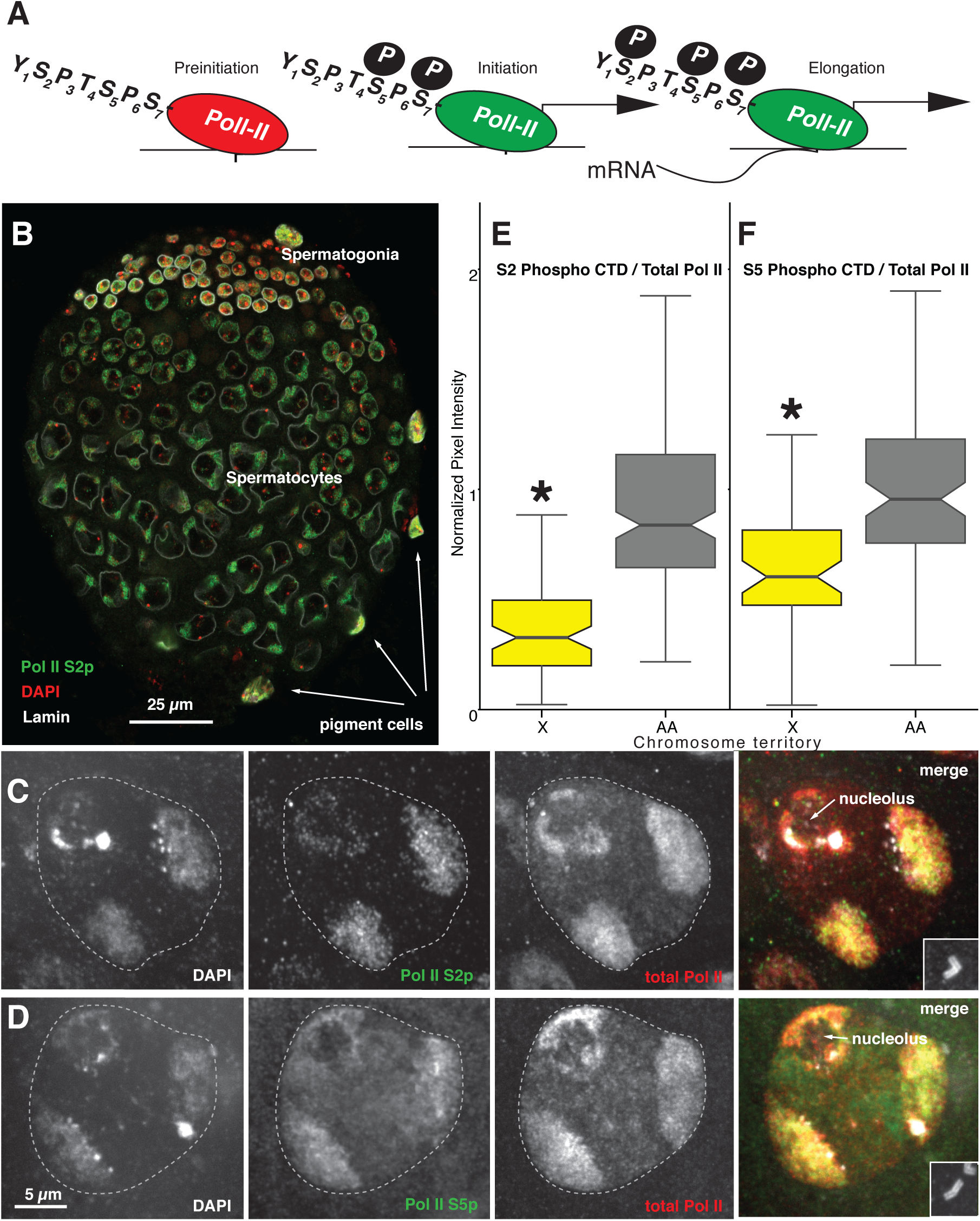
Territory-specific polymerase-II regulation. (**A**) Schematic of RNA Polymerase-II (Pol-II) with the C-terminal seven amino-acids (single letter code) in non-transcribing preinitiation (red), and initiation (green) stages at the promoter (bent arrow), as well as during elongation in the gene body. Key phosphorylation events (filled circles with “P”). (**B**) Third instar larval testis stained for Serine 2 phosphorylated (S2p) pol-II (green), nuclear lamin (white) and DAPI (red). Spermatogonia at the anterior (top) above spermatocytes are prominent. Polyploid pigment cells also shown (arrows). Scale is indicated. (**C-D**) Channel separated and merged images showing chromosome territories stained by DAPI (grey), total Pol-II (red), and (**C**) Pol-II S2p (green) or (**D**) Pol-II S5p (green). Scale indicated. Asterless stained centrioles from the same cell (inset). (**E-F**) boxplots (box = interquartile range (IQR), notch = 95% confidence interval of median, whiskers = ±1.5xIQR) showing staining intensities of active Pol-II relative to total Pol-II in X (yellow) and autosome pair (A/A, grey) territories. P-value ≤ 0.01, Welch’s t-test (*). (E) Pol-II S2p. (**D**) Pol-II S5p.

Our data clearly support a dynamic model, where X chromosomes are expressed at a higher rate in spermatogonia than one would expect based on DNA copy number alone, supporting the idea of at least partial X chromosome dosage compensation in the male germline. This is transient. We show that a major contributor to the unusual behavior of sex chromosomes in *Drosophila* germ cells is dramatic expression changes in the primary spermatocytes in G2 of meiosis-I. While lower expression of the X in early meiosis could be due to the absence of X-chromosome dosage compensation in the germline (*11*), and while the canonical dosage compensation pathway is absent in male germ cells (*22, 23*), there is evidence for non-canonical dosage compensation in testis (*20, 26*). Our data provides an important new argument against failed dosage compensation in the male germline, as the 4^th^ chromosome undergoes a similar dramatic decrease in steady-state transcript levels, despite being present in two copies. This suggests that sex chromosome, not copy number, determines activity in primary spermatocytes. Where X-like chromosomes are inactivated and the Y-like chromosomes are highly expressed.

The dynamic pattern of X chromosome expression is reminiscent of meiotic sex chromosome inactivation in other species. In mammals, the unusual pattern of X chromosome expression is similarly dynamic with high, possibly compensated, expression in spermatogonia, followed by X inactivation (*14*). It is possible that this is a special case of a more general inactivation of unpaired chromosome regions in a genomic defense model (*15*). This model is consistent with the inactivation of both the X and Y chromosomes in mammals. We observed two violations of the prediction that unpaired chromosomes are silenced in primary spermatocytes. Specifically, the 4^th^ chromosome, an ancient X chromosome that has re-acquired autosome status, would be active, and the Y would be inactive in the simplest versions of this model. While there are complicated models, where the pairing of the X and Y at the rDNA loci, or lack of pairing of the 4^th^ chromosome despite being present in two copies drive Y activation and 4^th^ inactivation, the most parsimonious hypothesis is that the evolved sex-chromosome nature of these chromosomes determines their activities. Thus, “sex chromosome nature” could be a conserved aspect of the regulation of chromosome wide gene expression in germ cells.

## Materials and Methods

### Sample preparation

We collected gonads from the immobile third instar (L3) larvae just prior to the final prepupal defecation and prior to eversion of the anterior spiracle (**Fig. S1A**). Staging was aided by staining gut contents using 0.5 mg/ml Sulforhodamine B yeast paste fed to L3 larvae (clear foregut, full midgut/hindgut) (**Fig. S1B**). We dissected L3 larval and 1-5 days old adult gonads and brains directly into Phosphate Buffered Saline, pH 7.4 (PBS). We used fixed gonads for immunostaining and gonads and brains for *in situ* hybridization (see the respective sections below).

For expression profiling whole gonads and single cells, initially we used enzymatic and mechanical treatments to remove non-gonadal tissue. We prepared frozen single-use aliquots of all enzymes used in preparing gonads/cells and tested to determine optimal digestion times. We removed fat body from gonads (**Fig. S1C-F**) using 150 U/µl Papain and 150 U/µl Collagenase in PBS at 22°C for 2-3 min in 1.5 ml Eppendorf tubes mixed by gently flicking the tube with a forefinger. We decanted floating fatbody cells and washed 2X with PBS. For whole gonad profiling, we saved the gonads in 350 μl RLT buffer mixed two times for 30 s, froze in a dry ice-ethanol bath and stored at -80 °C. We performed RNA-Seq on eight biological replicates each of cleaned L3 larval testes and ovaries. For single cell preparation, we pre-coated all tubes, pipettes, and filters with 0.5% Bovine Serum Albumin (BSA) to minimize loss of cells. We removed fat body from gonads as above, except we used 0.075% (w/v) Porcine Powdered Pancreas (PPP) rather than Papain (**Fig. S1C-D**). We dissociated gonads in 0.45% PPP and 750 U/µl Collagenase in PBS at 22°C for 30 min, teased with tungsten needles and pipetting under the dissecting scope. Digestion was stopped by adding Fetal Bovine Serum (FBS, 1% final v/v) for 2 min. We decanted the cell suspension and a wash of 0.04% BSA onto a 35µm cell filter. Cells were pelleted at 845 x g for 5 min and resuspended in 25 µl PBS, 0.04% BSA. A 1 μl cell suspension was used to calculate density microscopically. We performed scRNA-Seq on three biological replicates of L3 instar larval testes.

In initial trials, we determined that the somatic support cells that tightly encase the germline were separated and that the multicellular germline cysts were disrupted using gonads from flies expressing reporters (*tj>GFP* or *Vasa-GFP)* (**Fig. S1G-H**). We also stained cells with 1µg/ml Hoechst that detect nuclei and visualized small aggregates of cells (1%, with 4-8 cells, each 6 to 8μm), multinucleated cells (2%, mostly two nuclei spermatocyte cysts) and enucleated cells (0.5%) (n = 565 cells). We analyzed the viability of cells in 0.2% Trypan blue in 0.5X PBS, 0.02% BSA. 25-30% of cells took up dye, suggesting 70-75% viable cells prior to microfluidic loading (n=1000 cells) (**Fig. S1I**).

### Immunostaining

For immunostaining formaldehyde fixed tissues we used slightly different conditions depending on the antibodies used and location. For protein traps: we fixed in 4% formaldehyde in PBS for 20 min; blocked in 1X PBS plus 0.1% Triton X-100 (PBX), 3%BSA, 0.02% NaN_3_ and 2% Normal Goat Serum (NGS) for 30 min; incubated with primary antibodies (1:50 mouse α-Fas3, 1:50 mouse (1B1)hu-li tai shao, and 1:5000 chicken α-GFP in PBX) overnight at 4 °C; washed with 1X PBX; incubated with secondary antibodies (1:200 Alexa Fluor 488 α-chicken, 1:100 Alexa Fluor 568 α-mouse) with 1 µg/µl DAPI at room temperature for 2 hr; rinsed twice in 1X PBX, once in 1X PBS; and mounted onto a microscope slide in Vectashield. Images were acquired using Zeiss LSM 800 laser scanning confocal microscopes with 63x/1.4 Oil DIC M27x objective and illumination lasers at 405, 488, and 561 nm. For RNA Pol-II: we fixed in 9% formaldehyde in PBS for 20 mins; washed three times in PBS with 0.3% Triton X-100 (PBST); blocked for >1 hour in 5% NGS in PBST at room temperature; incubated in primary antibodies (1:1000 rat α-RNA polymerase II subunit B1, phosphorylated CTD Ser-2, or phosphorylated CTD Ser-5), 1:100 mouse α-lamin, 1:30,000 guinea pig α-Asl, 1:50 mouse α-Fas3, 1:10,000 chicken anti-GFP) in PBX with 5% NGS overnight at 4 °C; washed 3X with PBX for 10 min; incubated in secondary antibodies (1:1000 Alexa Fluor 488 goat α-rat, Alexa Fluor 568 goat α-mouse, and Alexa 647 goat α-guinea pig) at room temperature; counterstained with DAPI at 1:1000 in 5% NGS in PBX for 2-8 hr at room temperature; 3X in PBX for 10 min at room temperature and mounted in Aquapolymount under a No. 1.5 coverslip. Images of larval testes were acquired using Nikon Eclipse Ti with a 100X/1.4 NA oil immersion objective, a spinning disc confocal head, a CoolSNAP HQ2 camera and illumination lasers at 405, 491, 561, and 642 nm. The microscope was controlled by and images were acquired using MetaMorph. All data analysis was performed using *ImageJ*.

### in situ hybridization

To detect heterochromatin on X, 4^th^ and Y chromosomes, we synthesized probes (1.688 satellite of X chromosome and AATAT repeats for 4^th^ and AATAC repeats for Y chromosomes) using *D. melanogaster* genomic DNA as a template (*67, 73*). PCR was performed using 50-100 ng/μl of template DNA with Taq Platinum DNA polymerase following the manufacturer’s instructions. Probes were PCR-labeled with digoxigenin-11-dUTP or biotin-14-dUTP during their synthesis at the 5′ end. To detect the euchromatin, we synthesized oligopaint probes using 5’ fluorophore labeled and 5’ phosphorylated PCR primers, amplified with the following cycles: 95 °C for 5 min; 40 cycles of 95 °C for 30 s, 53 °C for 30 s and 72 °C for 15 s, with a final extension step at 72 °C for 5 min. The PCR product was purified using Zymo spin columns (D4031) and then digested using lambda exonuclease for 30 min at 37 °C and 10 min at 75 °C. The digested probe products were precipitated using ethanol and quantified using a nanodrop.

For mitotic metaphase chromosomes (**Fig. S5)**, we immersed dissected brains in 0.05% Colchicine in PBS for 20 min, hypotonized in tap water for 15 min, and fixed in 3:1 (v/v) ethanol:acetic acid for 15 min, transferred to 60% acetic acid and squashed on a slide. Fluorescence *in situ* hybridization for heterochromatic probes was performed according to Pinkel et al. (*74*) with modifications (*75*) using mitotic chromosome spreading or spermatocytes from adult testes. Post-hybridization washes were performed as follows: two times in 2X SSC at 42 °C for 5 min, two times in 0.1X SSC at 42 °C for 5 min, one time in 2X SCC at 42 °C for 5 min and finally in 2X SSC at room temperature for 10 min. Probes labeled with biotin-16-dUTP were detected using avidin-FITC conjugate (Fisher Scientific) and probes labeled with digoxigenin-11-dUTP were detected using anti-digoxigenin rhodamine (Roche). The preparations were counterstained using DAPI and mounted in Vectashield. Images from metaphase spreads and adult spermatocytes were obtained using a Zeiss Axiophot 2 microscope equipped with Bright field and epifluorescence optics.

For fluorescence *in situ* hybridization using oligopaints, we fixed testes in 5% formaldehyde washed in 1X PBX and then 0.3M NaCl, 30mM Na Citrate, 1% Tween 20 (2X SSCT), then successively in 20, 40, and 50% formamide in 2X SSCT. Testes were pre-denatured at 37 °C for 4 hr, 92 °C for 3 min, and 60 °C for 20 min in 50% formamide in 2X SSCT. We heated 100 pmol of the X-euchromatin primary probe and 200 pmol of the 2L-euchromatin primary probe in probe buffer with RNase A and testes to 91 °C for 3 min and incubated overnight at 37 °C for 18 hr on a shaker at 130 rpm. We washed testes in 50% formamide + 2X SSCT. Testes were then washed with decreasing concentrations of formamide (50 and 20%) and 2X SSCT. To counterstain with antibodies, we refixed with 16% Formaldehyde, 100% Tergitol, 10X PBS, heptane and MilliQ H20, washed in 1X PBX, and then blocked in a 1X PBX + 1.5% BSA mixture for 1 hr. Testes were incubated overnight at 4 °C with: 1:20 mouse α-lamin, 1:20 mouse α-lamin DmO, and 1:10,000 guinea pig α-Asl in PBX. Testes were washed in 1X PBX and then incubated in the secondary antibodies. Testes were mounted onto a microscope slide containing a drop of Vectashield. Images were acquired using Zeiss LSM 800 laser scanning confocal microscope with 63X/1.4 NA oil immersion objective and illumination lasers at 405, 488, 561, and 640 nm.

### Image analysis

To measure the X and 2L oligopaint territories, we loaded CZI images into *Imaris*. Centriole duplication is a marker of time during the prolonged G2 phase of primary spermatocyte development, which we scored using the Asterless (Asl) centriole marker. We created X and 2L oligopaint probe channel masks (n=23 for both X and 2L) of the pixel intensities in late primary spermatocyte cells. We measured volume and sphericity (shape) of the masks and performed Paired t-tests.

For quantitative measurements of Ser-2, Ser-5 Pol-II and total Pol-II signal, samples were dissected and processed in parallel for each experiment and imaged (at 16 bits) on a single day using identical microscope settings ensuring that all pixel intensities were within the dynamic range of the camera (no more than ¾ of the full dynamic range). We normalized X chromosome territory region of interest to the average total fluorescence intensity of the autosomes for each measurement in late apolar spermatocytes determined from that day’s experiment. The data presented are from two independent experiments. To quantify the amount of phosphorylated CTD Ser-2 or Ser-5, total Pol-II and DAPI in the discrete DNA domains within the nuclei of spermatocytes, autosomal or nucleolar associated DNA domains that were spatially separated from other domains (in XY and with no overlap in the Z-slices containing any signal from the cluster) were selected from images of spermatocytes. Z-slices encompassing the domain were then summed and the integrated pixel intensity for this volume was measured. An area of nucleoplasm not containing any other DNA domains was selected and measured for background subtraction. X chromosomal domain was identified by the nucleolus region and the other two were considered as autosomal domains. Ratios between X chromosomal and summed autosomal domains were calculated.

### RNA-Seq

For bulk sequencing, we added fatbody-free gonads to RNeasy 96 kit 350 μl RLT buffer mixed two times for 30 s, froze in a dry ice-ethanol bath and stored at -80 °C. We extracted total RNA from gonad samples using RNeasy 96 kit following the manufacturer’s protocol. We quantified RNA using the Quant-iT RiboGreen quantification kit and used 200 ng of total RNA for library preparation (*76*). We spiked in External RNA Control Consortium (ERCC) spike-ins pools 78A and 78B (transcribed from a certified standard reference) prior to fragmentation. For multiplexing, we used eight different TruSeq v2 kit barcoded adaptors. We sequenced stranded multiplexed libraries using a single-ended 50 bp strategy and generated RNA-Seq profiles of biological quadruplicates on a HiSeq Illumina 2500.

For single-cell sequencing, we loaded dissociated cells (6K for Replicates 1 and 2, 12K for replicate 3) onto the 10X Chromium system for barcoding and library preparation following the user guide for Single Cell 3’ Reagent Kits v2. We quantified libraries with Quant-iT PicoGreen and confirmed 300-500 bp insert sizes on a TapeStation 2200. We generated scRNA-Seq profiles in biological triplicate pools on the 10X Chromium System and sequenced (Read1 = 26 bp, Read2 = 98 bp and Readi7 = 8 bp) on a HiSeq Illumina 2500.

### RNA-Seq - Computational analysis

For the bulk RNA-Seq, we demultiplexed and converted Binary Base Call (BCL) files to FASTQ using bcl2fastq (v2.17.1.14, Illumina, San Diego, CA, USA). We processed FASTQ files through a custom RNA-Seq workflow (*77*); commit: 0609e5c8752). Briefly, reads are trimmed of Illumina adapters using *cutadapt* (*78*) with default parameter except for -q 20 and -- minimum-length=25. We mapped with *Hisat2* (*79*) with default parameters except --max-intronlen 300000 and --known-splicesite-infile using annotated splice sites (FlyBase r6-26). We removed multi-mapping reads and low-quality alignments using *samtools view* -q 20 (*80*). Gene expression and intergenic expression (FlyBase r6-26) is quantified in a strand-specific manner using *FeatureCount* from the *subread* package (*81*) with the -s 2 option (**Table S4**). The workflow outputs a number of quality control metrics from FastQC (*82*), FastQ Screen (*83*), Picard MarkDuplicates (*84*), and Picard CollectRnaSeqMetrics (*84*). In addition, we quantified intergenic expression and ERCC spike-in expression using *FeatureCount* (*85*). Bulk RNA-Seq data is available at the Gene Expression Omnibus GSE115478 (larval testes) and GSE115511 (larval ovaries). Bulk adult testes and ovaries are from a previously published data GSE99574 (*31*), **Table S1)**.

For scRNA-Seq, we converted BCL files to FASTQ using *cellranger mkfastq* (*86*). We demultiplexed cells, aligned reads, and quantified gene-level expression using *cellranger count* with default parameters (*86*). For alignment and quantification, we used the *D. melanogaster* genome assembly (dm6) with gene annotations from FlyBase r6-26. Larval testes scRNA-Seq data is available at the Gene Expression Omnibus GSE125947.

To quantify how well scRNA-Seq captured the overall expression patterns of the testis, we summed gene-level counts from all cells for each of our three replicates and normalized scRNA-Seq and bulk RNA-Seq libraries using Transcripts Per Million reads (TPM). We calculated pairwise Spearman rank correlation coefficients among all samples (**Table S1**).

Prior to the downstream analysis, we removed cell IDs that were likely to be empty and identified cell IDs that likely contain two or more cells known as multiplets (**Table S2**). There are two distinct classes of multiplets: homotypic and heterotypic. Homotypic multiplets contain 2 or more cells from the same cell-type. Homotypic multiplets tend to have high UMI counts and are found at the top of the UMI distribution however, setting an upper UMI threshold may bias against high RNA content cell-types. To identify an optimized upper UMI threshold, we performed a grid search over [4000, 5000, 6000] gene expression thresholds. For each point in the grid search, we clustered cells using *Seurat* (*87*). We then compared cluster calls using the adjusted rand index and selected an upper UMI threshold which did not drastically change clustering. We found that cluster calls remained stable using an upper threshold of ≤5,000 expressed genes, which removed 768 cells that were putative homotypic doublets. Heterotypic multiplets contain 2 or more cells from different cell-types. This leads to a mixture of expression profiles causing cells to look like intermediate cell-types, which are identifiable using *in silico* mixing of cell-types. We removed 410 cell IDs that behave like the mixture.

We combined all three scRNA-Seq replicates into one data set using single-cell integration (*88*) and clustered cells with K-Nearest Neighbors using the 2,000 most variably expressed genes (*87*). To determine cluster boundaries, we iteratively tested multiple threshold parameters (range 0.2-1.2), selecting a threshold of 0.3, which gives a set of 10 clusters that clearly separate germline and somatic cyst lineages, and each cluster is represented in all three replicates (**Table. S2**).

We compared gene expression patterns between scRNA-Seq and manually curated images (43 genes from the literature; 31 genes from this study; **Table S3**). We developed an overlap score (0-4) between scRNA-Seq expression and mRNA or protein expression in curated images. A score of 4 indicates a gene showed cell type biased expression (scRNA-Seq) in the exact same cell types as mRNA or protein expression (curated images). Since protein expression may lag transcription, we also gave a 4 when protein expression was later in the specific cell lineage. A score of 3 indicates gene was highly expressed, but not cell type biased, in the exact same cell types as mRNA or protein expression. A score of 2 indicates cell type biased gene expression in the same cell lineage as mRNA or protein expression. Finally, a score of 1 indicates high gene expression in the same cell lineage as mRNA or protein expression.

Expression by chromosome arms is complicated as gene expression of an individual cell is sparse in scRNA-Seq, thus simple aggregation of cell counts is not appropriate because missingness is not random (*89*) and could be confounded by chromosomal inactivation. Therefore, we compared X, 4^th^, and Y expression with the major autosomal arms (2^nd^ and 3^rd^) on an individual cell basis. We calculated the X:A,A, 4,4:A,A, and Y:A,A ratios for each cell by taking the total X-linked (or 4^th^-linked, Y-linked) reads divided by the total autosomal-linked reads (2^nd^ and 3^rd^) normalized by the number of genes per chromosome. We then permuted cell-type identity 10,000 times and calculated the number of times a the X:A,A, 4,4:A,A, or Y:A,A ratio were more extreme than observed values. To maximize completeness, we also performed permutation tests on a set of widely expressed genes (expressed in ≥ ⅓ of all cells; n = 589) which are effectively the genes expressed across multiple cell types.

We list reagents and resources used in our experiments in the FlyBase provided Author Reagent Table template (**Table S4**).

## Acknowledgments

This work utilized the computational resources of the NIH HPC Biowulf cluster (http://hpc.nih.gov), NCBI and FlyBase databases, and the Bloomington Drosophila Stock Center.

## Author contributions

SM, JMF, MDV, EM, and BO conceptualized the project; SM, JMF developed single cell methodology; JMF developed code; MA, MMLSP, CCA, DCCM, KC, ZD, KM, CAM, OMPG, ER, MS, and KY validated results; JMF, MA, BJG, CCA and CAM performed formal analysis; SM, MA, BJG, MMLSP, CCA, DCCM, KC, SD, ZD, KM, CAM, OMPG, ER, MS, KY, HES, and VS performed experiments; JMF curated metadata; SM, JMF, MDV, EM, and BO wrote the original draft; SM, JMF, MA, MDV, EM, and BO edited; SM, JMF, MA, CCA, MDV, and BO prepared figures; SM, JMF, HES, VS, NR, MDV, EM, and BO directed the work; SM and BO administered the project; NMR, MA, MDV, EM, and BO acquired funding.

## Data and materials availability

All data is available in the main text, the supplementary materials, and/or at NCBI Gene Expression Omnibus (GSE125947, GSE115511, and GSE115478).

