## Supplemental Figures for "Dynamic Sex Chromosome Expression in Drosophila Male Germ Cells"

**This PDF file includes:**

Figs. S1 to S6

**Other Supplementary Materials for this manuscript include the following:**

Table S1-S4

Movie1.

**Table S1. Bulk RNA-Seq of intact larval and adult gonads.** This table summarizes results related to bulk RNA-Seq in 3 worksheets. *Gene Level* contains raw gene level counts for bulk samples and gene annotations from FlyBase. *Spearman Among Samples* contains the complete Spearman correlation table comparing all bulk samples and scRNA-Seq samples. *Spearman With Clusters* is the complete Spearman correlation table but with scRNA-Seq samples split up into clusters.

**Table S2. Single cell RNA-Seq of third instar larval testis.** This table summarizes all results related to the single-cell RNA-Seq into 4 worksheets. *Cell Level Data* summarizes what 10x beads were selected as cells and the cell level X:A,A, Y:A,A, 4,4:A,A gene expression ratios. *Gene Level Data* summarizes gene level results including cluster level aggregated gene expression. *Cluster Level Data* summarizes the number of cells assigned to each cluster as well as the correlation among clusters. *One vs Rest* (*Biomarkers)* the complete differential expression table comparing each cluster with all other cells.

**Table S3. Genes used for annotation of single cell clusters.** To annotate clusters as putative cell types, we curated images from the literature and generated new cell type specific expression patterns using protein traps. (*20–50*)

**Table. S4. FlyBase Author's Resource Table.** This table provides additional details about stocks, reagents, software, and datasets used in this project. (*6*, *9*, *11–15*, *17*, *51–72*)

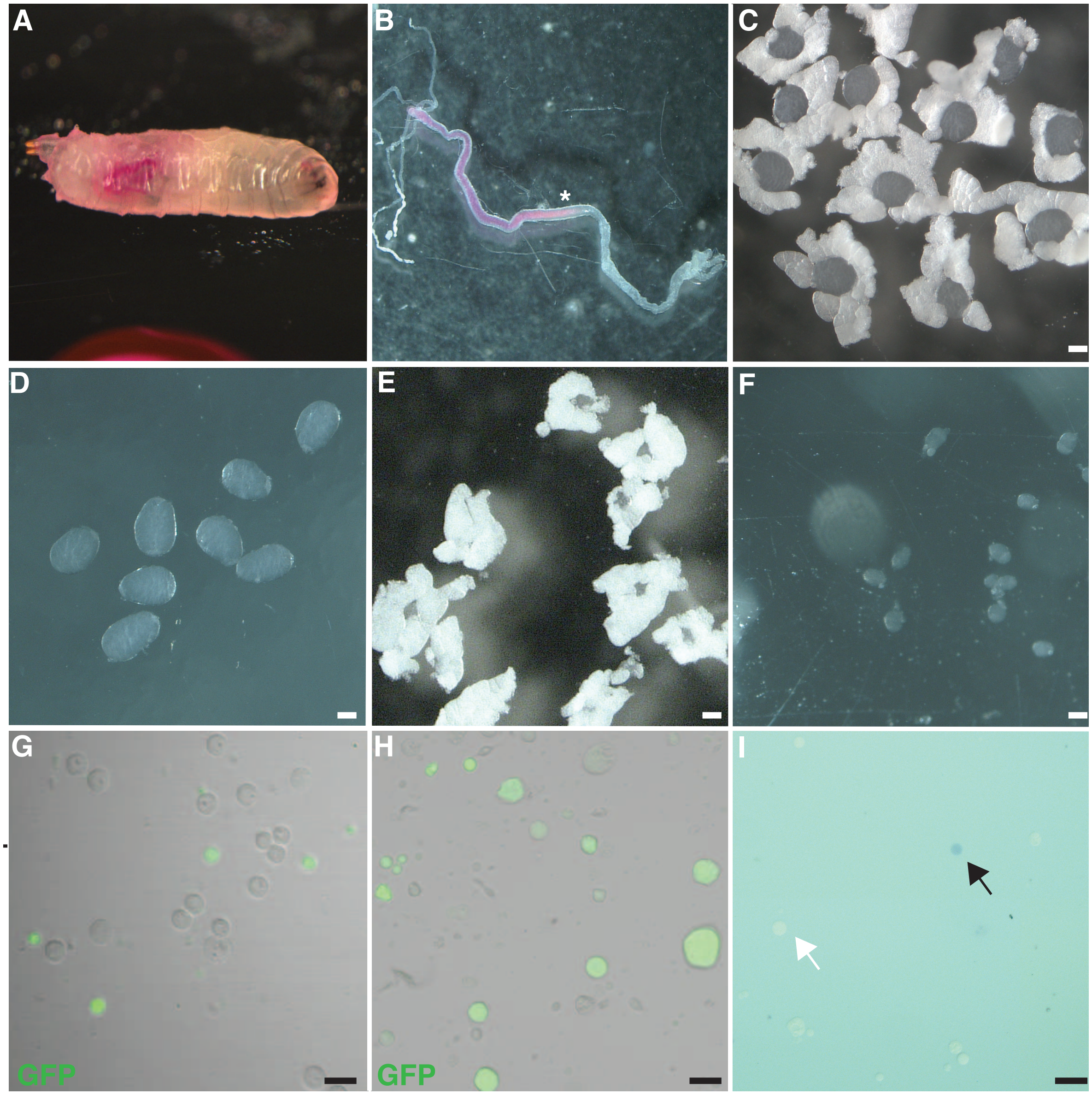


**Fig. S1**. **Single-cell preparation from Drosophila Larval testis**.

(**A**) Bright-field Microscopic image of late third instar larva fed with Sulforhodamine dye media (red). (**B**) Dissected larval gut with dye cleared from the foregut, and remaining in the midgut and hindgut but clear foregut. The boundary of food clearing in the midgut (Asterisk). (**C**, **E**) Dissected larval testes and ovaries with fat body. (**D**, **F**) Testes and ovaries cleaned of associated fatbody. Representative images (fluorescence and phase contrast) of dissociated cells marked for (**G**) *traffic jam* expression in the cyst cells (*tj-GAL4>UAS-GFP*) and (**H**) *vasa* expression in the germ cells (VASA-GFP), respectively. (**I**) Trypan blue staining indicates dead (black arrowhead) and live (white arrowhead) cells. White scale bars:100 µm, Black scale bars: 20 µm.

https://doi.org/10.35092/yhjc.11950746

**Fig. S2. UMAPS of all the genes.** For each gene we plot a UMAP project colored by Z-score value (-3, 3). The line graph represents normalized gene expression for each cluster.

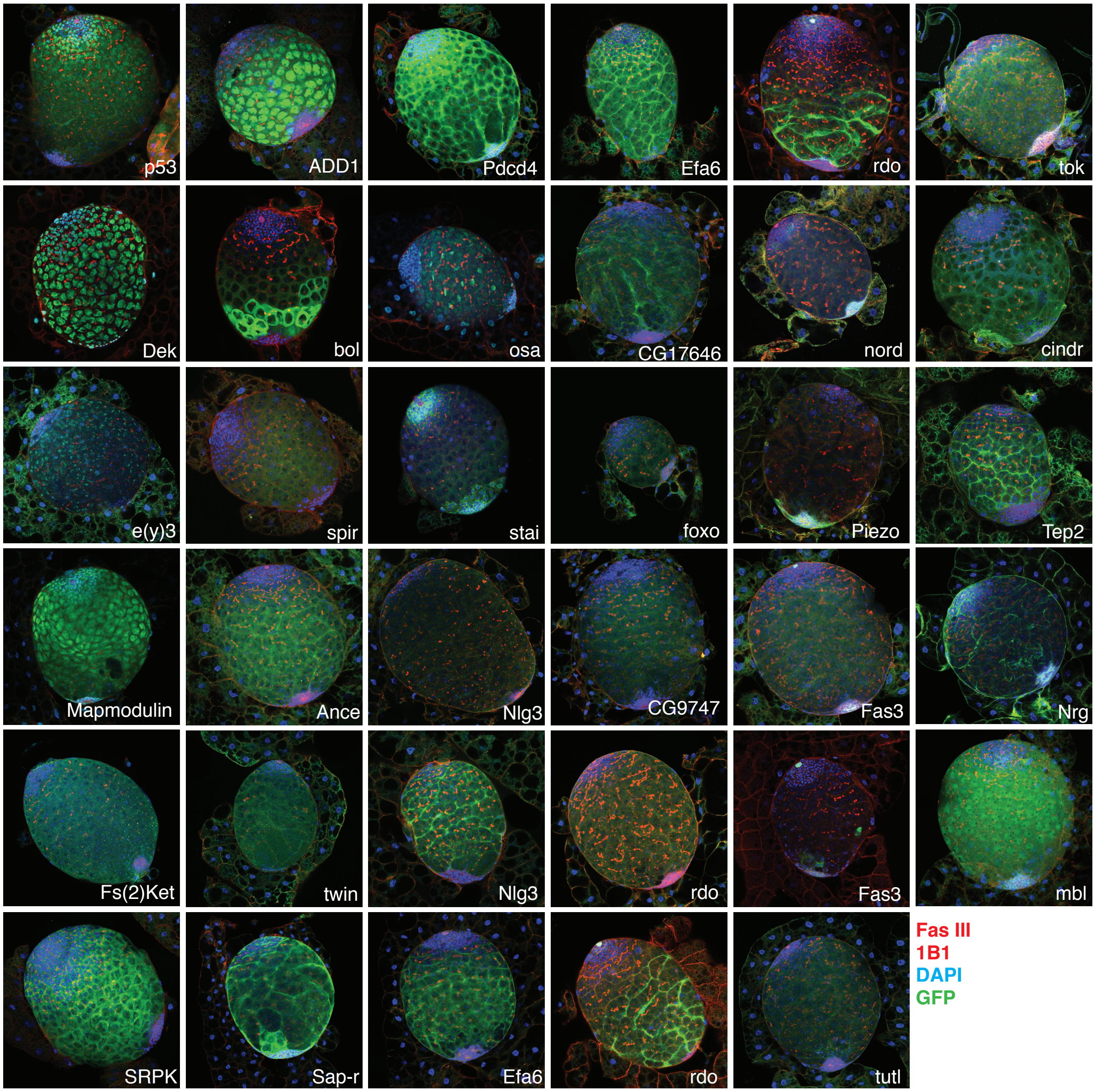


**Fig. S3. Immunofluorescence of protein traps used to annotate cell clusters from scRNA-Seq data.**

Immunofluorescence images showing protein expression of genes (green:GFP) from the protein trap reporters. Each panel consists of an individual gene, with color (Fas III, 1B1: red, DAPI: blue, and GFP: green).

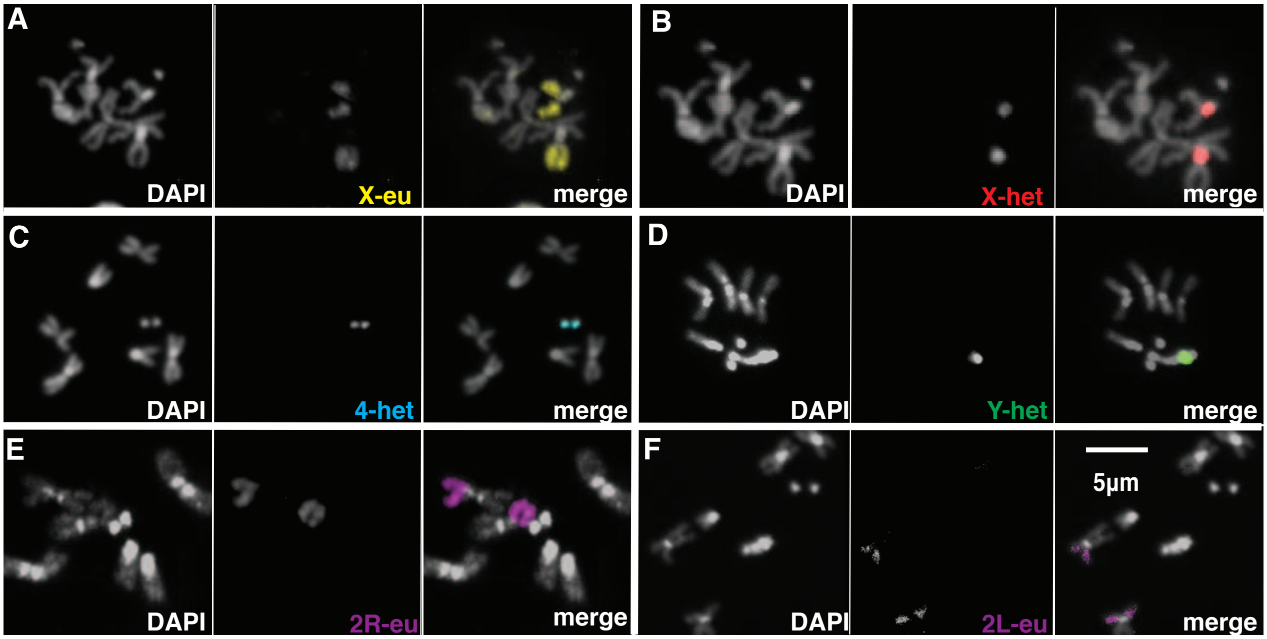


**Fig. S4. Spatial location of chromosomal territories in the Metaphase chromosome spreads from the brain.**

(**A**) Euchromatin oligopaints (yellow) detected a pair of telocentric chromosomes in the genome indicating X euchromatin territory (**B**) 1.688 satellite probe (red) detect the same telocentric chromosome in the same nucleus as in A, indicating the X heterochromatin (C)AATAT probe detected a pair of dot chromosomes indicating the 4th heterochromatin (**D**) AATAC probe detect the single chromosome indicating the Y heterochromatin. Euchromatin oligopaints (purple) detect the large metacentric chromosomes in (**E**) and (**F**) indicating either 2R or 2L euchromatin that is indistinguishable clearly. DAPI (grey).

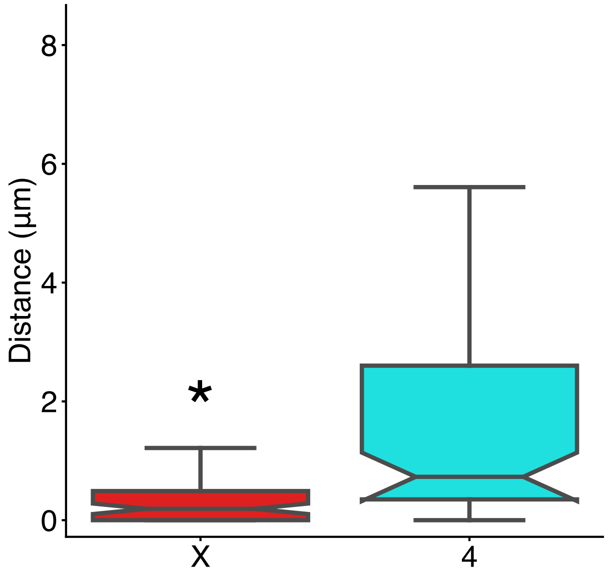


**Fig. S5. Relative distance of the X and 4th chromosomes to the nucleolus.**

Box plots showing the distributions of mean distances between the X (red), or 4th chromosomes (cyan), to the nucleolus. To measure chromosome distance to the nucleolus, we averaged the distance from the out heterochromatin probe edges to the closest nucleolus point. P-value ≤ 0.01 Wilcoxon signed-rank test (*).

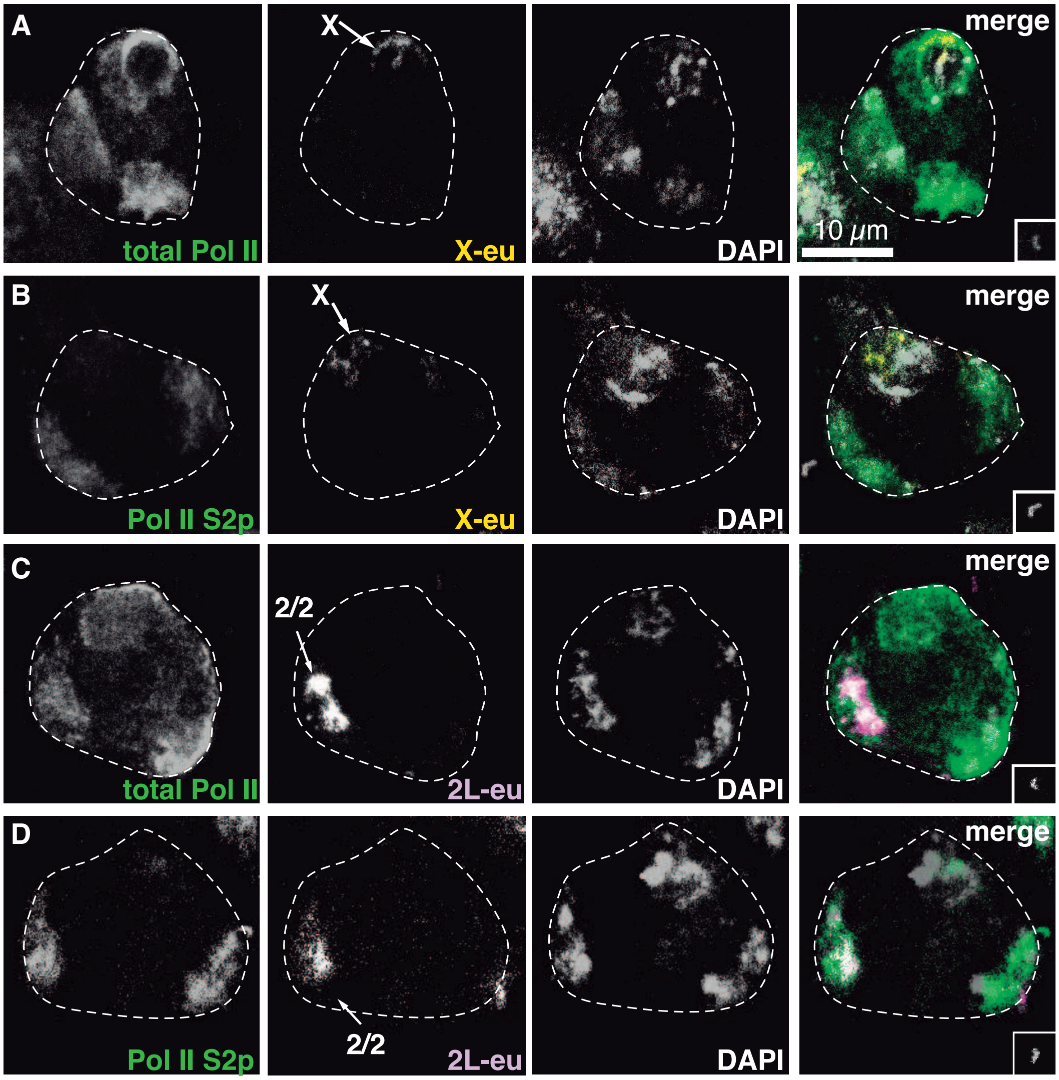


**Fig. S6. Localization of Total Pol-II and pSer2-CTD on X and 2L co-stained with euchromatic oligopaints**

Spatial localization of X chromosome euchromatin by oligopaints (yellow) with (**A**) total RNA Polymerase-II (total Pol-II) and (**B**) Serine 2 phosphorylated Pol-II (Pol-II S2p). Spatial localization of 2L chromosome euchromatin by oligopaints (purple) with (**C**) total Pol-II and (**D**) (Pol-II S2p). DAPI (grey), primary spermatocyte nucleus is outlined (white-dashed line) based on DAPI and bright-field images (not shown). Insets in the merged panels are Asterless stained centromere to stage individual cells.

**Movie.1. X and 2L spatial localization in primary spermatocytes by oligopaints.**

Spatial location of X chromosome euchromatin by oligopaints (yellow) and 2L chromosome euchromatin by oligopaints (purple) within a primary spermatocyte. DAPI (grey), Asterless stained centromere to stage individual cells (grey) and primary spermatocyte nucleus is outlined (white-dashed line) based on DAPI and bright-field images. Images are 'masked' in Imaris to obtain a 3D movie.

71. Cell Ranger. *10x Genomics*, (available at https://support.10xgenomics.com/single-cell-gene-expression/software/overview/welcome).

72. bcl2fastq. *Illumina, inc*, (available at https://support.illumina.com/sequencing/sequencing_software/bcl2fastq-conversion-software/downloads.html).
